# Radiocesium Uptake in Two Fungal Bed-cultivated Edible Mushroom Species in Forests in Fukushima, Japan

**DOI:** 10.64898/2026.01.28.702412

**Authors:** Masaru Sakai, Mirai Watanabe, Masami Kanao Koshikawa, Seiji Furukawa, Seiichi Takechi, Kaoru Yoshida, Akiko Takahashi, Masanori Tamaoki, Masabumi Komatsu, Hajime Murai, Takashi Tsuji, Mai Takagi, Seiji Hayashi

## Abstract

In Japan, mushrooms have long been valued as food resources, but the Fukushima nuclear accident disrupted production and shipment due to radiocesium contamination. Limited knowledge has particularly hindered outdoor fungal bed cultivation. To address this, we conducted cultivation experiments with *Lyophyllum decastes* and *Lepista nuda* across 14 broad-leaved deciduous forest sites in Fukushima Prefecture. Air dose rates at 1 m height ranged from 0.04 to 0.89 µSv/h. Radiocesium concentrations (combined ^137^Cs and ^134^Cs, expressed as Bq/kg at 90% water content) in fruit bodies were 0.4–12 (mean 2.3) for *Ly. decastes* and 0.2–43 (mean 8.1) for *Le. nuda*. Both species remained below Japan’s food safety threshold of 100 Bq/kg, indicating that safe cultivation is feasible across broad areas. Uptake patterns differed: concentrations in *Ly. decastes* correlated with contamination in litter and multiple soil layers, whereas in *Le. nuda* it correlated only with litter contamination. These findings suggest that clean soil fills for *Ly. decastes* and clean litter covers for *Le. nuda* could serve as mitigation strategies tailored to each species. Aggregated transfer factors (2.76 × 10^-5^ m^2^/kg for *Ly. decastes* and 6.60 × 10^-5^ m^2^/kg for *Le. nuda*) were lower than those reported for wild mushrooms. Overall, this study provides new insights into reducing radiocesium assimilation by cultivated mushrooms and supports the revival of outdoor fungal bed cultivation in contaminated landscapes.

## Introduction

Mushrooms collected from forests and cultivated outdoors, as both seasonal delicacies and sources of rural livelihood, have long been central to food culture [1–3]. However, radiocesium contamination has remained a serious challenge for mushroom cultivation and industry since the nuclear accidents such as in Chernobyl in 1986 [4,5] and in Fukushima in 2011 [6–8]. Fifteen years later, the shipment of wild mushrooms is still restricted in 55 of 59 municipalities of Fukushima Prefecture, due to ^137^Cs concentrations exceeding Japan’s food safety limit of 100 Bq/kg [9]. In addition, 17 municipalities continue to restrict shipments of log-cultivated mushrooms grown outdoors [9]. To revive mushroom cultivation and industry, science-based measures to produce less contaminated products are essential. A practical framework for addressing radiocesium contamination is to classify mushrooms into three groups—wild, log-cultivated, and fungal bed-cultivated—with the latter two further divided into outdoor and indoor systems.

Many species of edible wild mushroom accumulate large amounts of radiocesium, although the transfer factors vary widely among species and ecological guilds (e.g., mycorrhizal vs. saprophytic fungi) [6,7]. Soil improvements to mitigate radiocesium transfer are generally ineffective, as assimilation through extensive mycelial networks is too complex for localized interventions. Instead, scientific monitoring has identified species with lower transfer factors, which have been prioritized for collection, shipment, and consumption [6,7,10]. In fact, shipment restrictions for some of these low-accumulating species have been lifted in Japan since the Fukushima accident [11].

Among log-cultivated mushrooms, shiitake (*Lentinula edodes*) has been particularly affected, as Fukushima was one of Japan’s largest producers of that variety [12]. Radiocesium transfer from logs to mushrooms has been intensively studied [12–14], leading to government guidelines that have set a safe contamination threshold for logs (<50 Bq/kg) and have outlined cultivation management practices [15]. Based on this scientific guidance, log-cultivated mushroom production has gradually resumed in both indoor and outdoor settings in Fukushima [16].

Among fungal bed-cultivated mushrooms, indoor systems have resumed in Fukushima under official guidance [15], which instructs farmers to prepare less contaminated fungal beds (< 200 Bq/kg), similar to log-cultivated mushrooms. Meanwhile, radiocesium assimilation in outdoor fungal bed systems, which are directly exposed to natural soils and forest litter, remains poorly understood, hindering cultivation and underscoring the need for further research.

In this study, we examined radiocesium concentrations and environmental factors in two saprophytic mushroom species. Specifically, we investigated *Lyophyllum decastes* (wood/litter-decaying) and *Lepista nuda* (litter-decaying), using fungal bed systems at 14 broad-leaved deciduous forest sites in Fukushima, Japan. Both species are widely consumed in fried dishes and soups and had largely supported Fukushima’s mushroom industry before the accident. Wild *Ly. decastes* generally accumulate less radiocesium than other edible mushrooms, whereas *Le. nuda* tends to accumulate more [6]. Comparing assimilation patterns between these two species could therefore identify species-specific strategies for reducing contamination. We evaluated the environmental factors influencing radiocesium uptake, identified the soil layer serving as the primary source of nutrients, and assessed measures for obtaining less contaminated products. Our findings provide key insights for reviving outdoor fungal bed cultivation in contaminated landscapes.

## Methods

### Study sites and environments

The 14 experimental sites were in the villages of Iitate, Katsurao, Kawauchi, and Hirata; the towns of Minamiaizu, Shimogo, and Hanawa; and the cities of Koriyama (two sites), Kitakata, Soma, Date, Motomiya, and Tamura (Table S1). These are located in montane areas covered with broad-leaved deciduous forest dominated by Fagaceae species, typical of the Fukushima landscape. Field access did not require governmental permission because none of the sites were within the designated evacuation zone. Because the ultimate goal is to resume outdoor mushroom cultivation in inhabited zones, cultivation was not performed within the so-called ‘difficult-to-return’ zone, where ambient radiation levels remain relatively high. Approval was obtained from landowners at each site.

Cultivation beds of *Ly. decastes* and *Le. nuda* were planted at each site. The procedures for the cultivation experiments are summarized in Fig. 1. Prior to bed placement, ambient gamma radiation dose rates at 1 m above each plot (three plots per species per site) were measured with a scintillation survey meter (TCS 172B; Hitachi Aloka Medical, Tokyo, Japan) equipped with a NaI(Tl) probe. These readings were primarily attributable to ^137^Cs and primordial radionuclides in soils, although values may have been slightly elevated by background radiation and instrument noise. Differences among sites were interpreted as reflecting ^137^Cs accumulation in the soil, because airborne radiation levels are a reliable proxy for ^137^Cs soil inventories in contaminated areas [17,18]. These measurements are also publicly available for broad areas of Fukushima Prefecture via a national airborne monitoring website [19] and can be obtained with portable dosimeters accessible to the public.

**Fig. 1.**
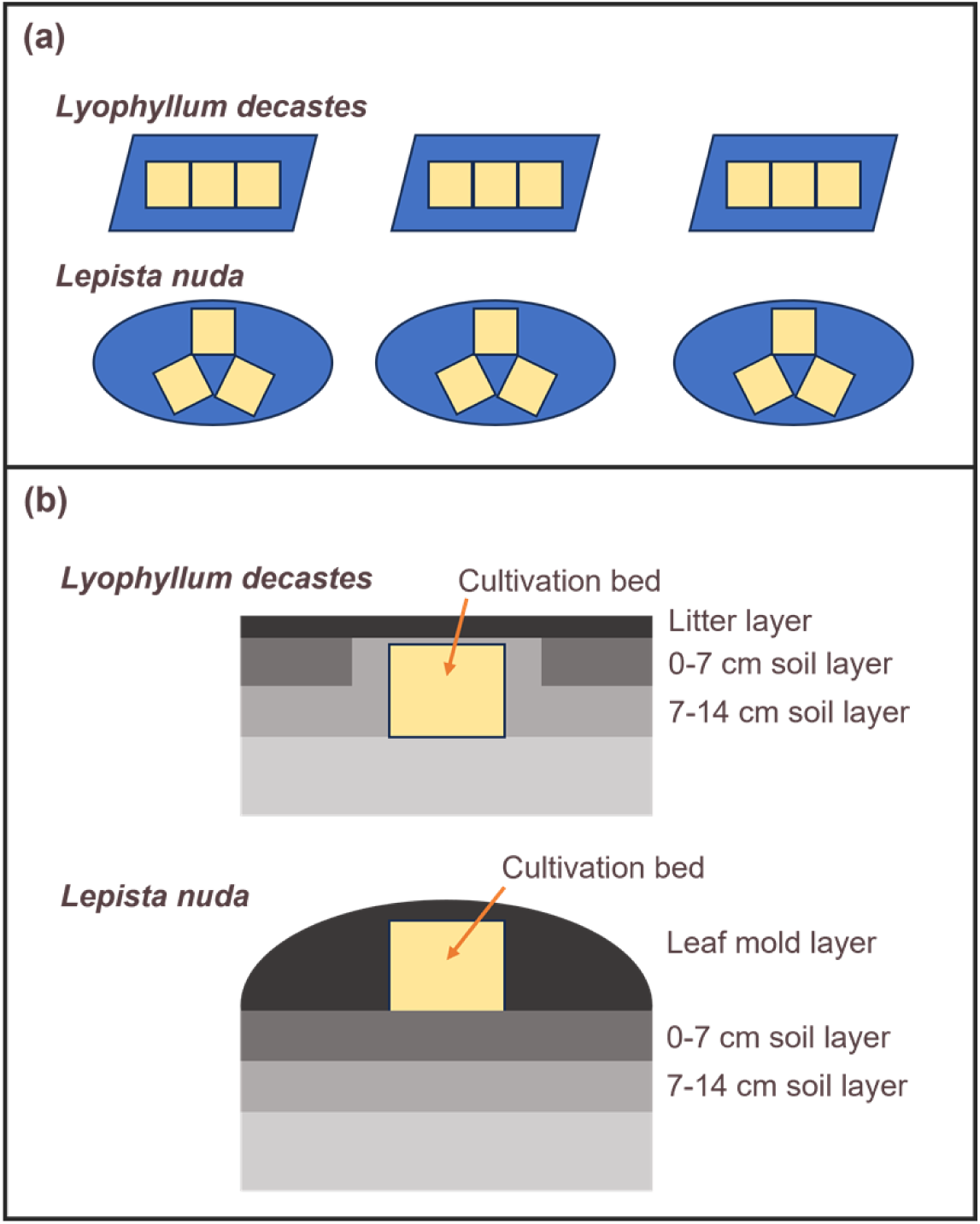
Layout and structure of cultivation beds for *Lyophyllum decastes* and *Lepista nuda* mushrooms in forests of Fukushima, Japan. (a) Top view of cultivation beds at the study site. (b) Cross-sectional view of cultivation beds and cover materials. Cultivation beds of *Ly. decastes* were placed on the top surface of soil of 14 cm depth. The sides and top of these beds were fully covered with 7–14 cm soil, which was further covered with litter. Cultivation beds of *Le. nuda* were placed on the topsoil and fully covered with leaf mold. To prevent wind disturbance of the leaf mold cover, wire nets and metal pegs were used to secure it. The distance between adjacent plots was consistent at 1 m across the six plots.

After measuring airborne radiation levels, the litter layer, the 0–7 cm soil layer (fermentation and humus layers plus surface mineral soil), and 7–14 cm soil layer (subsurface mineral soil) were separately collected from three plots of *Ly. decastes* at each study site (Fig. 1), because these strata contain most of the deposited ^137^Cs [20,21]. The total ^137^Cs soil inventory, as well as the inventories in the litter, 0–7 cm, and 7–14 cm layers (kBq/m^2^), were estimated from soil mass per unit area and ^137^Cs activity concentrations of the collected samples. The procedures for measuring ^137^Cs activity concentrations are described below. These inventory data were used as explanatory variables in statistical analyses for both species.

Meteorological conditions were monitored during cultivation experiments at each site (Table S1). Air temperature, relative humidity, and sunlight intensity were recorded at 10 min intervals using sensors mounted on 1.7 m poles near the cultivation beds (temperature and humidity: HOBO U23-001; Onset, Bourne, MA, USA; sunlight: HOBO UA-002-64; Onset). Temperature and humidity loggers were housed in white instrument shelters, while sunlight sensors were installed at the top of the poles. Mean values of air temperature and relative humidity, together with daily duration of sunlight exceeding 2,000 lux, were used as explanatory variables in subsequent statistical analyses.

### Fungal-bed cultivation

Three cuboid cultivation beds (12 × 20 × 13 cm) of *Ly. decastes* were placed in a hole (41 × 26 × 14 cm) created during soil sampling and fully covered with 7–14 cm soil, reflecting the species’ preference for buried woody substrate (Fig. 1). Then, the soil surface above each bed was covered with litter collected from the plots. In parallel, three cultivation beds of *Le. nuda* (12 × 20 × 13 cm) were placed on the mineral topsoil after removing the organic layers, and covered with leaf mold (litter mixed with fragmented organic material) collected from a 1 m diameter circle in each plot, consistent with its litter-decaying habit (Fig. 1). In total, 3 beds × 14 sites were established for each species.

The cultivation beds consisted of a fungus-inoculated 10:2 mixture of bark compost and bran. Because radiocesium concentrations in bark compost were low (∼50 Bq/kg) and bran was clean, radiocesium in cultivated mushrooms was assumed to originate primarily from surrounding litter and soils. A preliminary study also indicated that radiocesium concentrations in the fruit bodies of both species were < 1 Bq/kg when cultivated on the fungal bed substrate alone (MW, unpublished data). These beds were standard commercial products identical to those sold to farmers in Fukushima before the 2011 nuclear accident.

Beds were planted between 23 July and 1 August 2024. *Ly. decastes* was harvested in the period 7–20 October 2024, and *Le. nuda* in the period 7–21 November 2024. Cultivation periods ranged from 71 to 87 days for *Ly. decastes* and 102 to 118 days for *Le. nuda*, depending on growth. No water or fertilizers were supplied beyond natural rainfall. The total weight of mushrooms meeting shipping standards was recorded immediately after harvest.

### Radioactivity measurements

Litter and soil samples from *Ly. decastes* plots and leaf mold samples from *Le. nuda* plots were air-dried for at least 1 month. Litter and leaf mold were fragmented with scissors and sieved through 9.5 mm mesh, while soils were sieved through 2 mm mesh. Each sample was packed into a 100 mL plastic container. Dry weight was determined from the weight loss of subsamples dried at 105°C for ≥ 24 h, while bulk samples were stored at 25°C for other chemical analyses.

Fresh mushrooms were gently washed with tap water to remove adhering soil particles and organic debris, wiped, and dried at 60°C for ≥ 2 days. Drying at 60°C was chosen to minimize loss of volatile organic matter compared to drying at 105°C. Dried samples were finely ground with an electric mill and packed into 100 mL containers. Dry weights and sample densities were recorded prior to radioactivity measurements.

^137^Cs activity concentrations in litter, leaf mold, soil, and mushroom samples were determined via gamma-ray spectroscopy. Emissions at 661.6 keV (^137^Cs) were measured using a coaxial high-purity germanium detector (GC 2020; Canberra Japan, Tokyo, Japan). Because ^134^Cs from the Fukushima accident was below detection limits, ^134^Cs concentrations in mushrooms were estimated to obtain total ^137^Cs + ^134^Cs contamination levels [22]. We assumed that all ^137^Cs originated from the accident, with equal initial emissions of ^137^Cs and ^134^Cs. ^137^Cs activity concentrations in mushrooms on 11 March 2011 were estimated, substituted for ^134^Cs values at that date, and corrected to sampling dates using the ^134^Cs half-life (2.06 years).

Mean measurement accuracy was 5% for litter, leaf mold, and soil samples, and 8% for mushrooms (error counts/net area counts). Counting times ranged from 10,000 to 200,000 s. Activities were corrected for radioactive decay to the collection date. Mushroom concentrations were expressed on both dry weight and fresh weight bases, with fresh weight calculated assuming 90% water content [6,23].

The detector system was routinely calibrated with blanks (empty 100 mL containers) and ^137^Cs standards (CS401; Japan Radioisotope Association, Tokyo, Japan), and verified annually by the manufacturer using reference samples to ensure accuracy.

### Statistical analyses

To examine the relationship between total ^137^Cs soil inventory (kBq/km^2^) and airborne radiation levels at 1 m height (µSv/h), we constructed a linear mixed model (LMM). The response variable was log_10_-transformed total ^137^Cs soil inventory, the explanatory variable was log_10_-transformed radiation level, and study site identifiers were included as random intercepts to account for site variability. A Gaussian error structure was assumed. The resulting regression was used to estimate an upper threshold of airborne radiation at which combined ^137^Cs + ^134^Cs (hereafter, radiocesium) concentrations in mushrooms could exceed the Japanese food safety limit of 100 Bq/kg, contingent on statistically significant relationships between mushroom radiocesium concentrations and soil ^137^Cs inventories.

To evaluate factors influencing radiocesium concentrations in mushrooms, we constructed LMMs for each species. Explanatory variables included log_10_-transformed total ^137^Cs soil inventory (kBq/km^2^), mean relative humidity (%), total mushroom yield (g), cultivation period (days), mean daily duration of sunlight > 2,000 lux (h/day), and mean air temperature (°C). Multicollinearity was negligible (|*r*| < 0.7, variance inflation factor < 3). Soil inventories estimated in *Ly. decastes* plots were assigned to adjacent *Le. nuda* plots. Two models were compared: one using log_10_-transformed mushroom radiocesium concentrations (Gaussian family) and the other using raw concentrations (Gaussian family). The model with the lowest conditional Akaike information criterion (cAIC) was adopted for each species [24]. Site identifiers were included as random intercepts. Explanatory variables were standardized (mean = 0, standard deviation [SD] = 0.5) prior to analysis [25] to allow direct comparison of coefficients.

To identify major ^137^Cs sources, six additional LMMs were constructed, each with a single explanatory variable: log_10_-transformed ^137^Cs inventories or activity concentrations in litter, 0–7 cm soil, or 7–14 cm soil. For *Le. nuda*, an LMM with ^137^Cs activity concentration in leaf mold was also tested, given its likely role as a primary source. Site identifiers were included as random intercepts. All LMMs were fitted using the lmer function in the lme4 package [26]; explanatory variables were standardized with the arm package [27] in R 4.5.1 [28]. Statistical significance was set at *P* < 0.05 using the lmerTest package [29], and cAICs were calculated with the cAIC4 package [30].

Aggregated transfer factors for each species were calculated as the geometric mean ± geometric SD of radiocesium concentration in mushrooms divided by total ^137^Cs soil inventory (m^2^/kg), and compared to values reported in previous studies.

## Results

Air dose rates at 1 m height ranged from 0.04 to 0.89 µSv/h, and total ^137^Cs soil inventories ranged from 1.39 to 640 kBq/m^2^ across study sites. The LMM indicated a significant positive relationship between them (Fig. 2), expressed as:

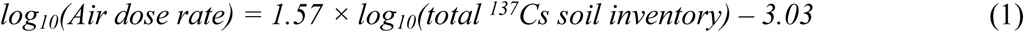

**Fig 2.**
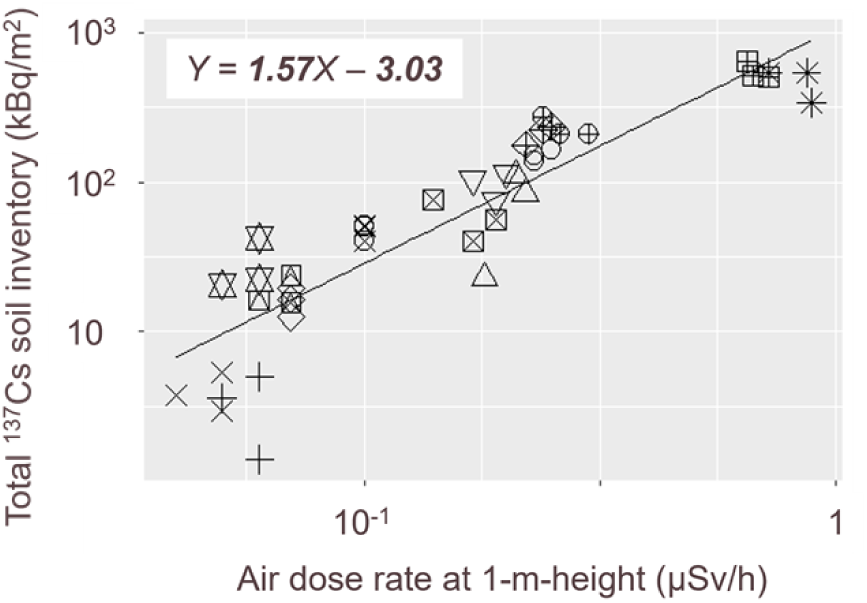
Relationship between the total ^137^Cs soil inventory and the air dose rate at 1 m height. Symbols represent individual study sites, with different shapes assigned to each site. The slope and intercept values were statistically significant (*P* < 0.05).

Radiocesium (^137^Cs + ^134^Cs) concentrations in mushrooms ranged from 0.4–12 Bq/kg (mean 2.3) in *Ly. decastes* and 0.2–43 Bq/kg (mean 8.1) in *Le. nuda* (90% water content basis). No samples exceeded the Japanese food safety limit of 100 Bq/kg. Conditional AICs for *Ly. decastes* were –49.8 (model 1, log_10_-transformed) and 122.1 (model 2, raw); for *Le. nuda* they were –22.9 and 185.6, respectively; thus, log_10_-transformed models were adopted for both species.

For *Ly. decastes*, radiocesium concentrations were significantly positively correlated with total ^137^Cs soil inventory and relative humidity (Table 1, Fig. 3). Based on equation (1), concentrations were estimated to exceed 100 Bq/kg at an air dose rate of 6.10 µSv/h, with other explanatory variables fixed at mean values. For *Le. nuda*, concentrations were significantly positively correlated with total ^137^Cs soil inventory (Table 2, Fig. 4), and exceedance was estimated at 5.91 µSv/h under mean conditions.

**Fig 3.**
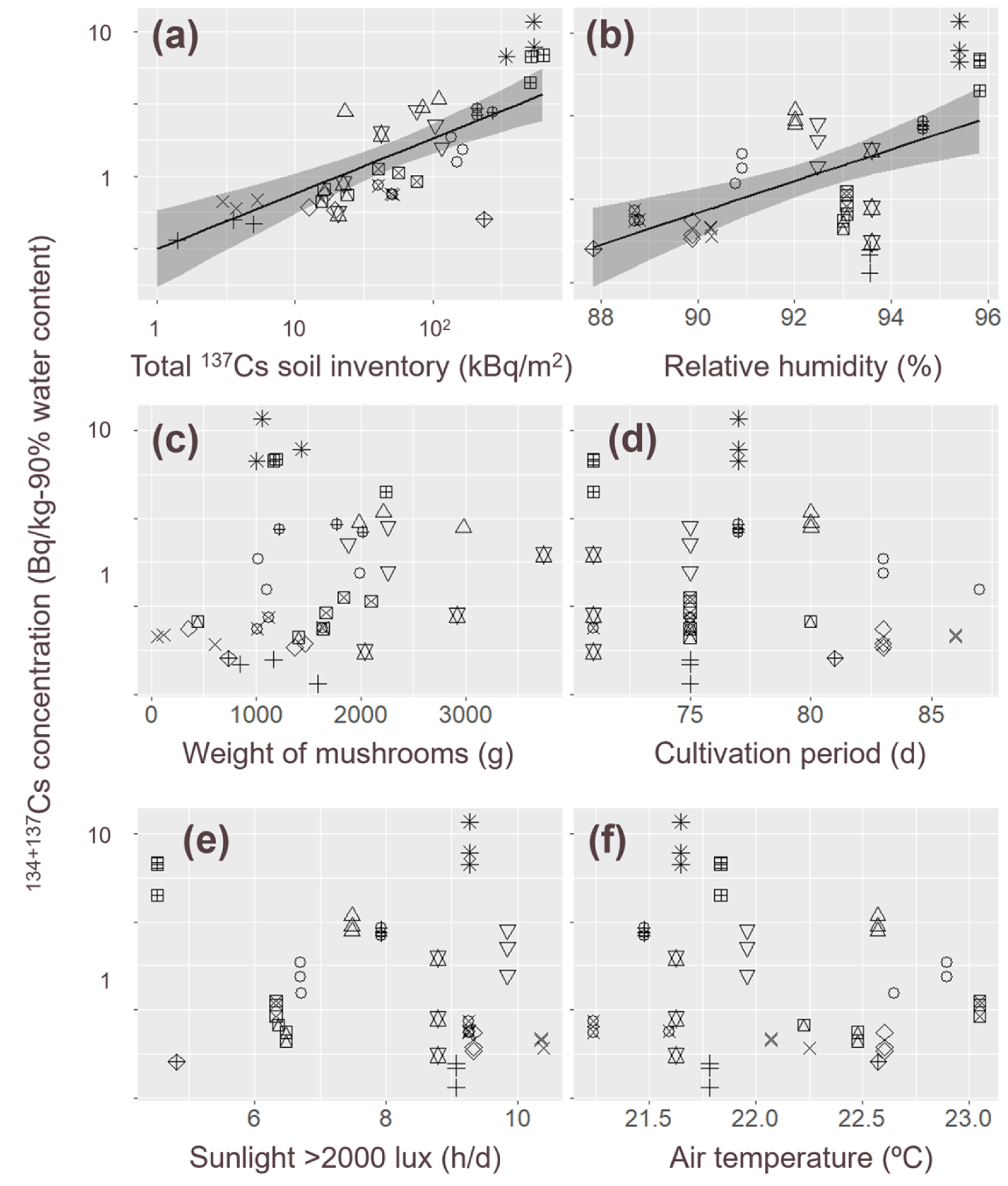
Relationships between the ^137^Cs + ^134^Cs activity concentration in *Lyophyllum decastes* mushrooms and environmental variables. (a) Total ^137^Cs soil inventory, (b) mean relative humidity, (c) total weight of harvested mushrooms, (d) cultivation period, (e) mean duration of sunlight > 2,000 lux, and (f) mean air temperature. Regression lines represent statistically significant relationships identified by the full linear mixed model, with other explanatory variables fixed at their mean values. Shaded bands indicate 95% confidence intervals. Different symbols denote individual study sites.

**Fig 4.**
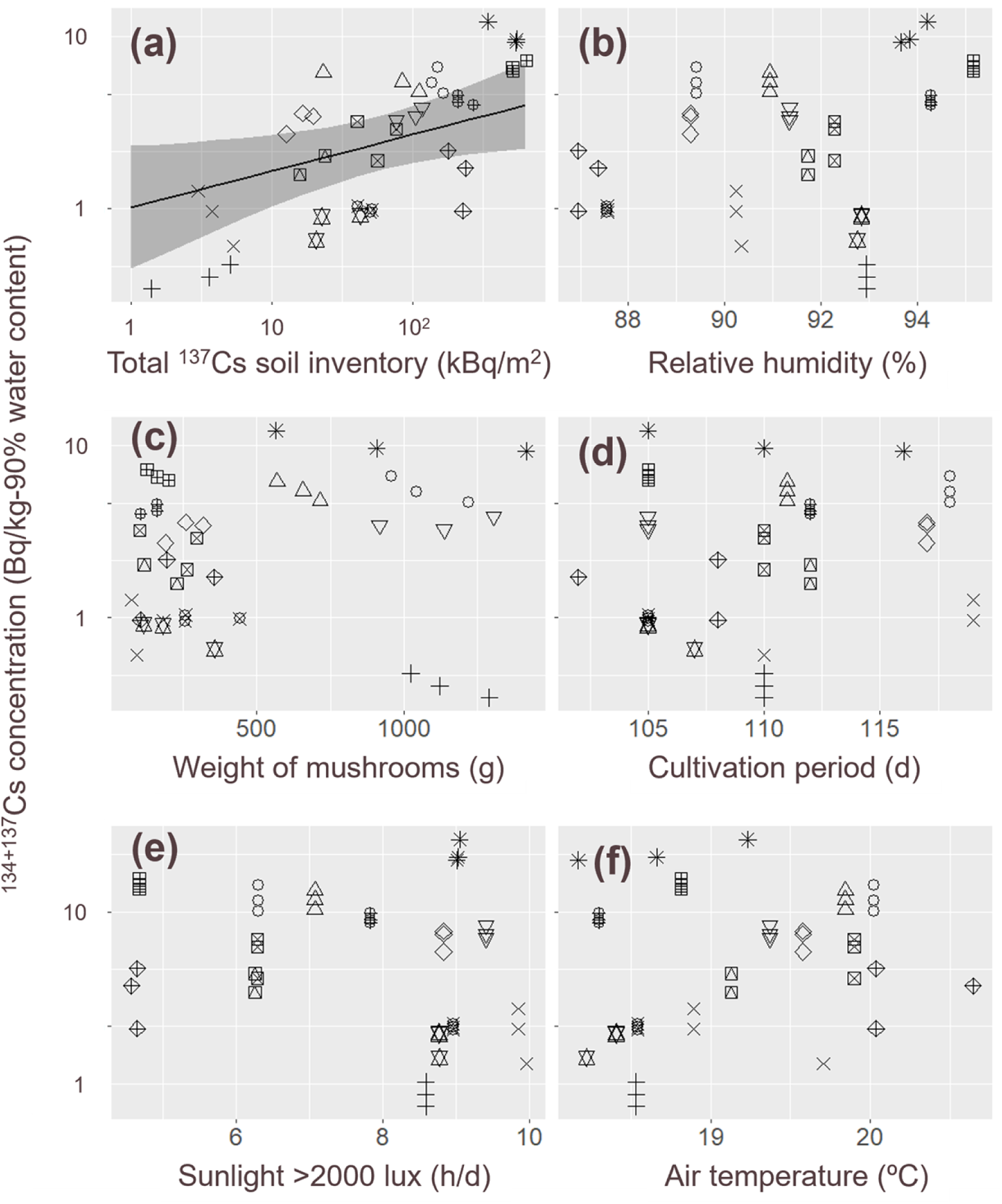
Relationships between the ^137^Cs + ^134^Cs activity concentration in *Lepista nuda* mushrooms and environmental variables. (a) Total ^137^Cs soil inventory in soil, (b) mean relative humidity, (c) total weight of harvested mushrooms, (d) cultivation period, (e) mean duration of sunlight > 2,000 lux, and (f) mean air temperature. Regression lines represent statistically significant relationships identified by the full linear mixed model, with other explanatory variables fixed at their mean values. Shaded bands indicate 95% confidence intervals. Different symbols denote individual study sites.

**Table 1.** Results of a linear mixed model testing the effects of total ^137^Cs inventory in soil, mean relative humidity, total weight of harvested mushrooms, cultivation period, mean duration of sunlight > 2,000 lux, and mean air temperature on ^137^Cs + ^134^Cs activity concentrations in *Lyophyllum decastes* mushrooms. Statistically significant values are shown in bold.

|  | Estimate | SE | *t* | *P* |
| --- | --- | --- | --- | --- |
| Total $^{137}\text{Cs}$ inventory in soil | 0.54 | 0.10 | 5.43 | **0.00** |
| Relative humidity | 0.42 | 0.11 | 3.97 | **0.00** |
| Cultivation period | 0.17 | 0.10 | 1.63 | 0.11 |
| Sunlight > 2000 lux | 0.17 | 0.12 | 1.48 | 0.16 |
| Weight of mushroom | 0.08 | 0.06 | 1.30 | 0.20 |
| Air temperature | -0.01 | 0.11 | -0.05 | 0.96 |

**Table 2.** Results of a linear mixed model testing the effects of total ^137^Cs inventory in soil, mean relative humidity, total weight of harvested mushrooms, cultivation period, mean duration of sunlight > 2,000 lux, and mean air temperature on ^137^Cs + ^134^Cs activity concentrations in *Lepista nuda* mushrooms. Statistically significant values are shown in bold.

|  | Estimate | SE | *t* | *P* |
| --- | --- | --- | --- | --- |
| Total $^{137}\text{Cs}$ inventory in soil | 0.45 | 0.19 | 2.36 | **0.03** |
| Relative humidity | 0.62 | 0.27 | 2.30 | 0.10 |
| Weight of mushroom | −0.18 | 0.14 | −1.26 | 0.22 |
| Cultivation period | 0.46 | 0.17 | 2.67 | 0.05 |
| Sunlight > 2000 lux | 0.04 | 0.27 | 0.14 | 0.90 |
| Air temperature | 0.48 | 0.23 | 2.09 | 0.11 |

Species-specific source relationships were evident (Figs. 5, 6). For *Ly. decastes*, ^137^Cs inventories in 0–7 cm and 7–14 cm soils, and ^137^Cs activity concentrations in litter and both soil layers were significantly positively correlated with mushroom radiocesium concentrations (Table 3, Fig. 5). In *Le. nuda*, the ^137^Cs inventory and activity concentration in litter, and the activity concentration in leaf mold, were significantly positively correlated (Table 3, Fig. 6).

**Fig 5.**
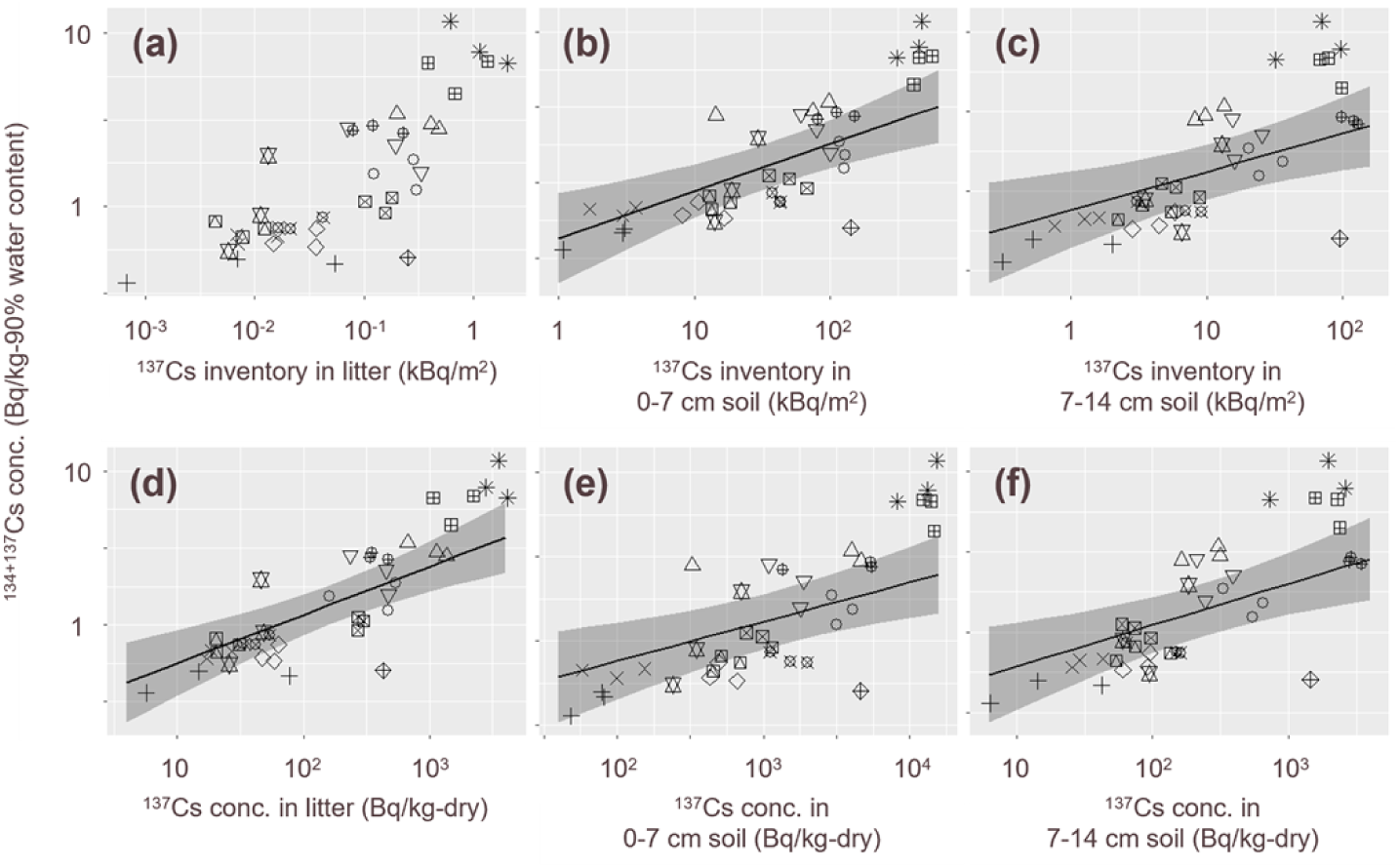
Relationships between the ^137^Cs + ^134^Cs activity concentration in *Lyophyllum decastes* mushrooms and the ^137^Cs inventory and concentration in litter and soils. ^137^Cs inventories in (a) litter, (b) 0–7 cm soil, and (c) 7–14 cm soil. ^137^Cs activity concentrations in (d) litter, (e) 0–7 cm soil, and (f) 7–14 cm soil. Regression lines represent statistically significant relationships identified by linear mixed models. Shaded bands indicate 95% confidence intervals. Different symbols denote individual study sites.

**Fig. 6.**
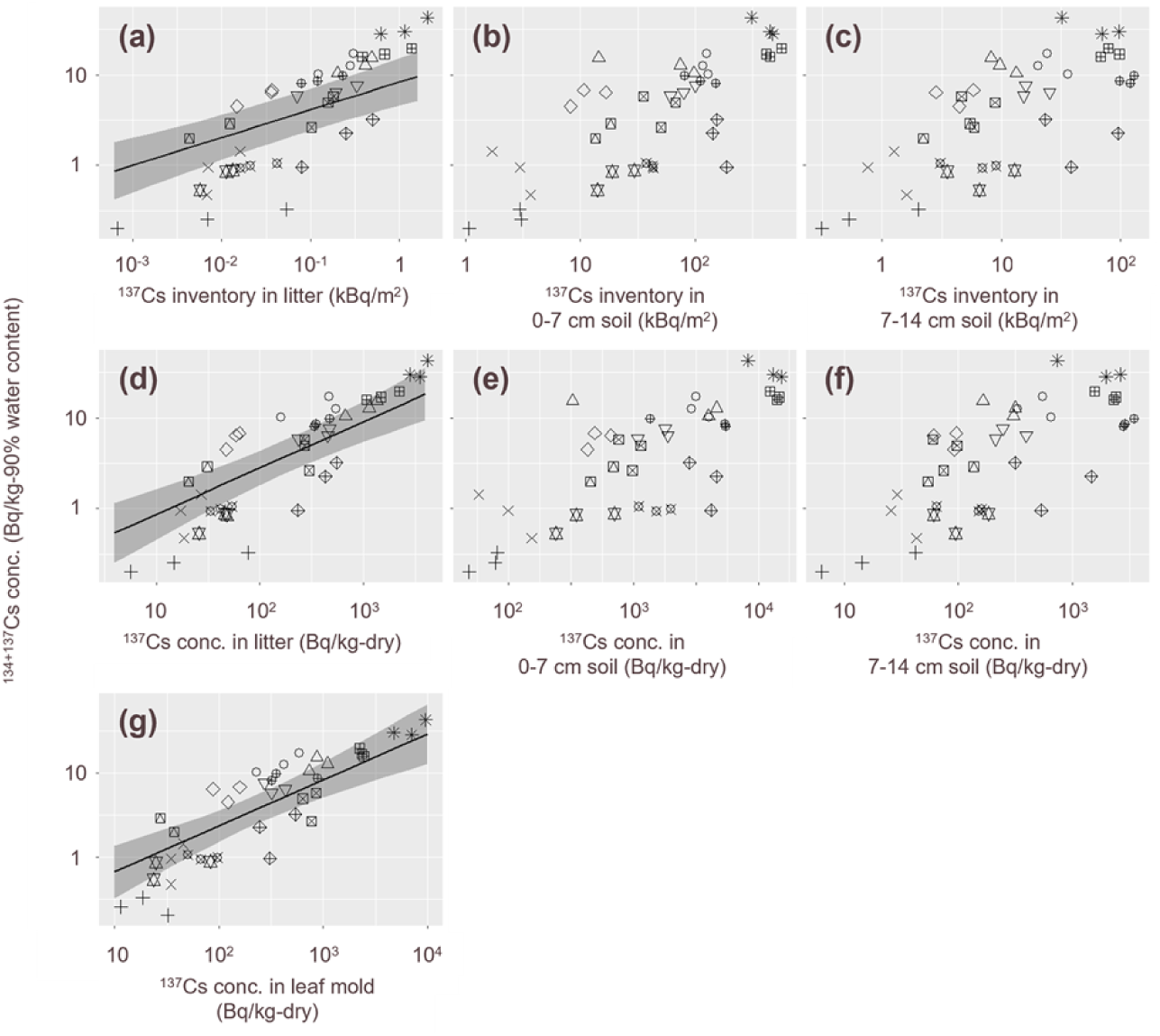
Relationships between ^137^Cs + ^134^Cs activity concentrations in *Lepista nuda* mushrooms and ^137^Cs inventories and activity concentrations in litter, soil, and leaf mold. ^137^Cs inventories in (a) litter, (b) 0–7 cm soil, and (c) 7–14 cm soil. ^137^Cs activity concentrations in (d) litter, (e) 0–7 cm soil, (f) 7–14 cm soil, and (g) leaf mold. Regression lines represent statistically significant relationships identified by linear mixed models. Shaded bands indicate 95% confidence intervals. Different symbols denote individual study sites.

**Table 3.** Results of 12 linear mixed models testing the effects of ^137^Cs inventory and concentration in litter, 0–7 cm soil, and 7–14 cm soil on ^137^Cs + ^134^Cs activity concentrations in *Lyophyllum decastes* and *Lepista nuda* mushrooms. Statistically significant values are shown in bold.

|  | Estimate | SE | *t* | *P* |
| --- | --- | --- | --- | --- |
| *Lyophyllum decastes* |  |  |  |  |
| $^{137}\text{Cs}$ inventory in litter | 0.10 | 0.06 | 1.74 | 0.09 |
| $^{137}\text{Cs}$ inventory in 0–7-cm-depth soil | 0.31 | 0.08 | 3.96 | **0.00** |
| $^{137}\text{Cs}$ inventory in 7–14-cm-depth soil | 0.24 | 0.08 | 3.03 | **0.00** |
| $^{137}\text{Cs}$ concentration in litter | 0.31 | 0.07 | 4.60 | **0.00** |
| $^{137}\text{Cs}$ concentration in 0–7-cm-depth soil | 0.23 | 0.07 | 3.28 | **0.00** |
| $^{137}\text{Cs}$ concentration in 7–14-cm-depth soil | 0.26 | 0.08 | 3.28 | **0.00** |
| *Lepista nuda* |  |  |  |  |
| $^{137}\text{Cs}$ inventory in litter | 0.31 | 0.05 | 5.97 | **0.00** |
| $^{137}\text{Cs}$ inventory in 0–7-cm-depth soil | 0.18 | 0.12 | 1.47 | 0.15 |
| $^{137}\text{Cs}$ inventory in 7–14-cm-depth soil | 0.15 | 0.11 | 1.37 | 0.18 |
| $^{137}\text{Cs}$ concentration in litter | 0.51 | 0.08 | 6.29 | **0.00** |
| $^{137}\text{Cs}$ concentration in 0–7-cm-depth soil | 0.01 | 0.10 | 0.13 | 0.90 |
| $^{137}\text{Cs}$ concentration in 7–14-cm-depth soil | 0.15 | 0.11 | 1.34 | 0.19 |
| $^{137}\text{Cs}$ concentration in leaf mold | 0.55 | 0.10 | 5.75 | **0.00** |

The aggregated transfer factors were 2.76 × 10^-5^ ± 1.17 m^2^/kg for *Ly. Decastes* and 6.60 × 10^-5^ ± 1.18 m^2^/kg for *Le. nuda*.

## Discussion

This study confirms that radiocesium activity concentrations in *Ly. decastes* and *Le. nuda* mushrooms cultivated in broad-leaved deciduous forests are well below the Japanese food safety limit of 100 Bq/kg. Model estimates indicated that concentrations would reach 100 Bq/kg at 6.10 µSv/h for *Ly. decastes* and 5.91 µSv/h for *Le. nuda*. Such elevated levels are currently restricted to the ‘difficult-to-return’ zones in Fukushima [19]. These findings suggest that both species generally exhibit low radiocesium accumulation when cultivated outside the evacuation zone near the Fukushima Daiichi nuclear power plant. Moreover, the results provide practical insights into measures for mitigating radiocesium assimilation in cultivated mushrooms.

For *Ly. decastes*, radiocesium concentrations were consistently low (0.4–12 Bq/kg) but showed a significant positive correlation with the total ^137^Cs soil inventory, indicating slight assimilation from soils proportional to contamination levels. Similar positive relationships have been reported in certain wild vegetable species [18,31], underscoring that cultivation outside high-radiation areas is a prerequisite for safe crop production. Relative humidity also exhibited a positive correlation with mushroom radiocesium concentrations. Because soil moisture strongly influences radiocesium bioavailability [32] and fungal uptake [33], higher humidity may indirectly increase assimilation in *Ly. decastes*.

For *Ly. decastes*, ^137^Cs inventories in 0–7 cm and 7–14 cm soils, as well as ^137^Cs activity concentrations in litter and both soil layers, were positively correlated with mushroom radiocesium concentrations. These results suggest assimilation from both litter and multiple soil horizons. Our data indicate that ^137^Cs was primarily stored in 0–7 cm soil (mean 77% of total), followed by 7–14 cm soil (mean 23%), and litter (mean 0.23%), consistent with previous studies [20,21]. In contrast, radiocesium bioavailability is generally higher in organic layers and decreases with depth [34,35]. This balance likely explains the observed transfers to *Ly. decastes* from multiple layers. Because the cultivation beds were placed in holes that were filled with 7–14 cm soil, and the soil surface above each bed was covered with litter (Fig. 1), the cultivation beds were in direct contact with 7–14 cm soil. Meanwhile, the surrounding forest floor retained natural soil profile, consisting of litter layer overlying 0–7 cm and 7–14 cm soils. Thus, although the cultivation beds directly contacted 7–14 cm soil, they could also be indirectly influenced by radiocesium present in the adjacent litter, 0–7 cm soil, and 7–14 cm soil. Consequently, *Ly. decastes* could assimilate radiocesium from all layers. Practically, filling cultivation bed holes with less contaminated soil, as done here, appears to be an effective mitigation measure. This should be verified through manipulation experiments using soil fills with varying contamination levels.

For *Le. nuda*, radiocesium concentrations in mushrooms (0.2–43 Bq/kg) remained below national safety limits for shipment and consumption. As with *Ly. decastes*, the total ^137^Cs soil inventory was positively correlated with mushroom contamination, suggesting that cultivation at less contaminated sites is a promising strategy. When focusing on direct relationships, radiocesium concentrations in *Le. nuda* mushrooms were higher at sites with greater ^137^Cs in litter and leaf mold. Because *Le. nuda* is a litter-decaying fungus that extends mycelia into surrounding litter and produces fruiting bodies from it [36,37], these results clearly confirm that assimilation occurs primarily from litter. Although radiocesium concentrations in mushroom were low at our study sites, maintaining clean litter cover during cultivation is an effective mitigation measure. This should be tested through manipulation experiments using leaf mold covers with varying contamination levels.

Our results confirm that *Ly. decastes* and *Le. nuda* mushrooms contain radiocesium concentrations low enough to meet the national safety limit for shipment and consumption, and are broadly obtainable throughout broad-leaved deciduous forests with < 1 µSv/h. The geometric mean of the aggregated transfer factor was one order of magnitude lower in this study for *Ly. decastes* (2.6 × 10^-5^ m^2^/kg) than for previously reported wild *Ly. decastes* mushrooms (1.0 × 10^-4^ m^2^/kg in 2011–2017 [6] and 4.0 × 10^-4^ m^2^/kg in 2016–2020 [38]). For *Le. nuda*, the value was two orders lower (6.6 × 10^-5^ m^2^/kg) than those of wild *Le. nuda* mushrooms (2.4 × 10^-3^ m^2^/kg in 2011–2017 [6] and 2.8 × 10^-3^ m^2^/kg in 2016–2020 [38]). These results suggest that both temporal changes and cultivation environments (wild versus managed beds) are critical factors determining radiocesium contamination in these mushroom species, and highlight the potential of outdoor cultivation to provide less contaminated products compared to wild mushroom collections.

Our findings further indicate that filling cultivation bed holes with clean soil for *Ly. decastes* and preparing clean leaf mold cover for *Le. nuda* effectively reduces contamination in cultivated mushrooms. Nonetheless, our study had limitations that warrant future research. First, the thresholds at which *Ly. decastes* (6.10 µSv/h) and *Le. nuda* (5.91 µSv/h) mushrooms exceed 100 Bq/kg was simply estimated based on our low dose rate sites (0.04–0.89 µSv/h), and thus uncertainty still remains on the estimations. For example, highly contaminated sites may still provide radiocesium not only from litter and soil but also from canopy layers to mushrooms [39,40]. This implies that radiocesium concentrations in mushrooms may reach at 100 Bq/kg in lower dose rate sites than expected. Therefore, additional experiments at more contaminated sites are necessary when we want to know the actual thresholds. Second, it was confined to broad-leaved deciduous forests, and different assimilation patterns may occur in other forest types such as evergreen coniferous forests [39,40]. Broader assessments across multiple forest types are needed. Third, soil adhering to mushrooms, which was removed in this study, can substantially increase measured radiocesium concentrations. This should be considered in future work. Finally, we did not quantify the relative contributions of surrounding natural materials (litter and soil) and fungal bed substrates to mushroom contamination. To establish robust guidelines for bed preparation and cultivation practices, it is important to do so.

To date, limited knowledge of radiocesium transfers to edible mushrooms cultivated outdoors has hindered the resumption of field production of mushrooms in contaminated areas. The present study provides the first evidence that *Ly. decastes* and *Le. nuda* can be successfully cultivated across a broad range of contamination levels in broad-leaved deciduous forests. Science-based approaches of this kind are crucial to revive local mushroom culture and industry in polluted regions. Expanding research to other edible mushroom species is essential to advance contamination management strategies in outdoor fungal bed cultivation.

## Acknowledgments

We thank the landowners who allowed us to conduct cultivation experiments.

## Supporting information

S1 Table. Characteristics of radiocesium assimilation by two fungal bed-cultivated edible mushroom species (*Lyophyllum decastes* and *Lepista nuda*) in forests of Fukushima, Japan. The table shows the study sites, raw data on ^137^Cs + ^134^Cs activity concentrations in each species, and the environmental variables. (XLSX)

## Author contributions

**Conceptualization:** Mirai Watanabe, Seiji Hayashi, Seiji Furukawa, Masami Kanao Koshikawa, Masaru Sakai

**Data curation:** Masaru Sakai, Mirai Watanabe, Masami K. Koshikawa, Akiko Takahashi, Seiichi Takechi, Kaoru Yoshida, Seiji Furukawa

**Formal analysis:** Masaru Sakai, Mirai Watanabe

**Funding acquisition:** Seiji Hayashi, Mirai Watanabe

**Investigation:** All authors

**Methodology:** Masaru Sakai, Mirai Watanabe, Masami Kanao Koshikawa, Seiji Furukawa, Seiji Hayashi, Masabumi Komatsu, Masanori Tamaoki

**Visualization:** Masaru Sakai, Mirai Watanabe

**Writing – original draft:** Masaru Sakai

**Writing – review & editing:** All authors

